# Needlestack: an ultra-sensitive variant caller for multi-sample next generation sequencing data

**DOI:** 10.1101/639377

**Authors:** Tiffany M. Delhomme, Patrice H. Avogbe, Aurélie Gabriel, Nicolas Alcala, Noemie Leblay, Catherine Voegele, Maxime Vallée, Priscilia Chopard, Amélie Chabrier, Behnoush Abedi-Ardekani, Valérie Gaborieau, Ivana Holcatova, Vladimir Janout, Lenka Foretová, Sasa Milosavljevic, David Zaridze, Anush Mukeriya, Elisabeth Brambilla, Paul Brennan, Ghislaine Scelo, Lynnette Fernandez-Cuesta, Graham Byrnes, Florence Le Calvez-Kelm, James D. McKay, Matthieu Foll

**Affiliations:** Genetic Cancer Susceptibility Group, Section of Genetics, International Agency for Research on Cancer (IARC-WHO), 150 cours Albert Thomas, 69008, Lyon, France; Genetic Epidemiology Group, Section of Genetics, International Agency for Research on Cancer (IARC-WHO), 150 cours Albert Thomas, 69008, Lyon, France; Institute of Hygiene and Epidemiology, Charles University, 1st Faculty of Medicine, Prague, Czech Republic; Faculty of Health Sciences, Palacky University, Olomouc, Czech Republic; Department of Cancer Epidemiology and Genetics, Masaryk Memorial Cancer Institute, Brno, Czech Republic; International Organization for Cancer Prevention and Research (IOCPR), Belgrade, Serbia; Russian N.N. Blokhin Cancer Research Centre, Moscow, The Russian Federation; Centre Hospitalier Universitaire de Grenoble Département d’Anatomie et Cytologie Pathologiques, CS 10217 Grenoble, France; Section of Environment and Radiation, International Agency for Research on Cancer (IARC-WHO), 150 cours Albert Thomas, 69008, Lyon, France

## Abstract

The emergence of Next-Generation Sequencing (NGS) has revolutionized the way of reaching a genome sequence, with the promise of potentially providing a comprehensive characterization of DNA variations. Nevertheless, detecting somatic mutations is still a difficult problem, in particular when trying to identify low abundance mutations such as subclonal mutations, tumour-derived alterations in body fluids or somatic mutations from histological normal tissue. The main challenge is to precisely distinguish between sequencing artefacts and true mutations, particularly when the latter are so rare they reach similar abundance levels as artefacts. Here, we present needlestack, a highly sensitive variant caller, which directly learns from the data the level of systematic sequencing errors to accurately call mutations. Needlestack is based on the idea that the sequencing error rate can be dynamically estimated from analyzing multiple samples together. We show that the sequencing error rate varies across alterations, illustrating the need to precisely estimate it. We evaluate the performance of needlestack for various types of variations, and we show that needlestack is robust among positions and outperforms existing state-of-the-art method for low abundance mutations. Needlestack, along with its source code is freely available on the GitHub plateform: https://github.com/IARCbioinfo/needlestack.

## INTRODUCTION

Massive parallel sequencing, or next generation sequencing (NGS), has revolutionized the manner in which genetic variation can be explored, due to a large increase in throughput and unprecedented ability to detect low-abundance variations compared to the traditional Sanger sequencing, and at a greatly reduced cost per sequenced base. However, because these new technologies are prone to errors, identifying genetic variants from NGS data remains a considerable challenge (1). This is particularly true in heterogeneous samples, where the variant allelic fractions (VAF, the ratio of the number of sequencing reads carrying the mutant allele to the total read count) deviate away from the expectations of a diploid genome (0%, 50% or 100% for the three possible diploid genotypes), until the point where the mutant alleles make up only a small fraction of the sequenced reads, approaching the background error rate. Nevertheless, robustly identifying low VAF sequence variants in such heterogeneous settings can be highly informative, for example providing insights into the clonal evolution of tumours (2), analyzing the cell-free DNA in order to identify tumour-derived molecular footprints (3), or evaluating somatic mutations in histologically normal material (4).

The error rate of next generation sequencing is known to vary across DNA base pairs and even across multiple base changes at the same DNA position (5,6). NGS errors originate from many of the steps in the sequencing process, stemming from the quality of the template DNA, its subsequent fragmentation, the library preparation, the base calling, or the alignment step subsequent to the sequencing of raw reads. Some of these errors have a tendency to reoccur consistently across samples whereas others have a more unpredictable appearance. The net effect of NGS being made of errors from multiple sources is that they become highly difficult to distinguish or correct (7). Variant identification methods that consider this highly variable error pattern may improve our ability to robustly detect true sequence variants even when their abundance is low. Most current algorithms use a probabilistic model on VAF applied independently across samples to distinguish between sequencing artifacts and true variations (8), while methods specifically designed to detect low abundance mutation, like shearwaterML (9,10), propose to benefit from the shared knowledge on errors across samples, but are limited by the requirement of a prior threshold on the error rate.

Here, we have explored the approach of using multiple samples analyzed concurrently to develop an error model for each potential base change. Sequence variants are identified as outliers relative to this robust error model. This method, called needlestack, allows the identification of sequencing variants in a dynamic manner relative to the variable error pattern found in NGS data, and is particularly appropriate to call variants that are rare in the sequenced material. By combining this method with additional laboratory processing for further error correction (11) and very deep next generation sequencing, we are able to robustly identify VAFs well below 1% while maintaining acceptable false discovery rates. We conducted multiple rigorous performance estimations and comparisons with methods for both somatic and germline variant detection. We deployed our pipeline focusing on efficiency and robustness using the Domain-Specific Language (DSL) nextflow (12), and on reproducibility by providing Docker and Singularity images. Source code is versioned and freely available on GitHub (https://github.com/IARCbioinfo/needlestack).

## MATERIAL AND METHODS

### Needlestack overview

Needlestack estimates for each candidate alteration, *i.e*. each pair of position and base change (the three non-reference nucleotides and each observed insertions and deletions) the systematic sequencing error rate across a series of samples, typically more than twenty to ensure a reasonable estimation of this metric. Then, for each sample, it computes the *p*-value for the observed reads under the null hypothesis of this estimated model of errors, and transforms this *p*-value into a Phred-scale Q-value reported as a variant quality score (QVAL) for the candidate mutation. As such it measures the evidence that the observed mutation is not explained by the error model, and should therefore be considered a mutation.

Needlestack takes as input a series of BAM files, and is based on three main piped processes, the generation of the mpileup file containing read counts at the target positions using samtools (13), the reformatting of this file into readable tabulated file and finally the estimation of the error model using our R regression script (see below) coupled with the computations of Q-values (Supplementary Figure 1). Needlestack is highly parallelizable as input positions are analyzed independently. As an output, needlestack provides a multi-sample VCF file containing all candidate variants that obtain a QVAL higher than the input threshold in at least one sample, general information about the variant in the INFO field (*e.g*. error rate estimation, maximum observed QVAL) and individual information in the GENOTYPE field (*e.g*. QVAL of the sample, coverage of the sample at the position).

### The Needlestack algorithm

Let *i=1…N* be the index of the sample taken from an aligned sequenced panel of size *N, j* the genomic position considered and *k* the potential alteration, with *k* ∈ (A,T,C,G,ins,del), *ins* and *del* covering respectively every insertion and deletion observed in the data at position *j*. Let DP_ij_ denote the total number of sequenced reads at position *j* for the sample *i*, AO_ijk_ the reads count supporting alteration *k* and *e*_jk_ the corresponding error rate. We model the sequencing error distribution using a negative binomial (NB) regression (without intercept):

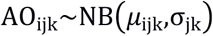

with *σ*_kj_ the over-dispersion parameter and *μ*_ijk_=e_jk_ * DP_ijk_ corresponding to the expected number of reads supporting alteration *k* across samples with a coverage *D*_ijk_. A robust negative binomial regression method (14) is employed to ensure that the outliers from this error model, such as true mutations, are not biasing the regression parameters estimates. This model is based on a robust weighted maximum likelihood estimator (MLE) for the over-dispersion parameter *σ*_jk_. We modified the original implementation of this regression to fit the need of our model here with: (*i*) a linear link function, (*ii*) a zero intercept, as a null coverage will exhibit a null read count, and (*iii*) an approximation of the bounding functions to allow the MLE to run efficiently for high coverage data (see Supplementary Methods).

For each position *j* and alternative *k*, we perform this robust negative binomial regression to estimate parameters *σ*_jk_ and *σ*_kj_. We then consider a sample *i* as carrying a true mutation *k* at the position *j* when being an outlier from the corresponding error model. We calculate for each sample a *p*-value for being an outlier using the estimated parameters that we further transform into *q*-values using the Benjamini and Hochberg procedure (15) to account for multiple testing and control the false discovery rate.

Importantly, because true mutations are identified as the outliers from the error model fitted using a robust regression, this approach is more suited to detect low-abundance mutations. Common mutations (for example germline SNPs with common allele frequencies) will be observed in the error model and therefore not detected as outliers by needlestack. In practice we found that mutations with a minor allele frequency below 10% can be accurately detected (see below). Additionally, while allowing over-dispersion, our model assumes that the error rate *σ*_jk_ is homogeneous across samples for a given alteration. This means that it should be applied to a homogeneous series of samples (that is prepared using comparable laboratory techniques and sequencing machines etc.). Importantly other types of errors that have less tendency to reoccur uniformly across samples are identified by needlestack as outliers.

### Sequencing data for performance evaluation

125 cell-free DNA (cfDNA) samples from healthy donors were used to study the distribution of error rates estimated by needlestack and to estimate its accuracy to detect low VAF using *in-silico* mutations. We also sequenced 46 cfDNA samples from 18 small-cell lung cancer (SCLC) patients and 28 squamous-cell carcinoma (SCC) patients, two cancer types that harbour a high prevalence of *TP53* mutations (respectively 99% (16) and 81% (17)). In order to validate in the tumour the low VAF mutations identified by needlestack in the cfDNA, we also sequenced tumour samples for these patients. Each of the cfDNA samples was sequenced for the *TP53* exonic regions (exons 2-11, which corresponds to 1,704 base pairs with a median coverage of around 10,000X) using the IonTorrent Proton technology, in two technical independent duplicates in order to account for potential errors during library preparation. Details about cfDNA sequencing steps and tumour sequencing method are provided in the Supplementary Material.

Additionally, we performed whole-exome sequencing (WES) from the blood of 62 samples from an independent cohort in order to estimate the performance of needlestack on germline mutations. As a gold standard, we used genotypes derived from Illumina SNP array (Illumina 5M beadarray) that were available for 33 of these 62 samples.

### Comparison with other variant callers

We used BAMsurgeon software (18) to introduce single nucleotide variations (SNVs) at varied VAF in the 125 cfDNA samples in *TP53* in order to benchmark and compare the method through *in-silico* simulations. BAMsurgeon presents the advantage of synthetic benchmarking methods that allow the simulation of mutations for which gold standards don’t exist to evaluate the performance (here low VAF, that are in addition challenging to validate), while maintaining the real data background such as the true error profiles. We introduced 1000 SNVs at random positions in the gene in random samples, and we replicated this process in ten batches. As each sample has been sequenced twice, we introduced each *in-silico* mutation in the two technical duplicates of a sample. We took benefit from the variable coverage among samples and genomic positions to study the sensitivity of our method down to VAF=10^−4^. For each mutation *m*, the VAF was simulated using a log-uniform distribution: VAF_*m*_=10^−*u*^ with *u*~uniform(0,4). Mutations were only introduced at positions where at least five mutated reads would be observed. This means that a mutation with a VAF=10^−4^ would be introduced only in positions with a coverage of at least 50,000X. To compare needlestack with a similar variant caller, we ran ShearwaterML (4,10) on the same ten batches (see Supplementary Methods). ShearwaterML is based on a beta-binomial regression and requires an *a-priori* threshold *t* for the error rate. ShearwaterML excludes each sample having a number of alternative bases higher than t*coverage, aiming at removing potential true mutations that act as outliers in the regression to robustly estimate the error rate. To compute the global performance of both methods, the ten simulation batches were merged, and only mutations detected in both technical duplicates were considered. In-silico simulations were repeated for 1-base pair insertions and deletions (indels) for needlestack. In this case, the total number of *in-silico* mutations was reduced to minimize the potential alignment artifacts created by the introduction of two indels close together. For that, using the same initial data, 100 insertions and 100 deletions were added again in ten simulations batches (total of twenty batches).

To estimate the ability of needlestack to detect rare germline variations, we used the 62 WES from blood samples. Needlestack variant calling was performed using our germline recommendations (see Supplementary Methods). GATK variant calling was performed using HaplotypeCaller best practice workflow (19) (see Supplementary Methods). From the 3,446,898 bead array non-reference genotypes (0/1 or 1/1) distributed over 113,232 positions in the 33 individuals, we selected 20,439 genotypes with a sufficient coverage (see Supplementary Methods). In a second part, to account for possible bias in the array, variant calls from both needlestack and GATK were compared independently of the array data, on a total of 44,314,972 exonic positions. To compare only rare germline variants, we removed common variants from each calling set (bead array, GATK calling and needlestack calling, see Supplementary Methods).

### Error rate estimation

To estimate the error rate variability across positions, we computed with needlestack the sequencing error rates from two data sets of the *TP53* gene sequenced with two different technologies (on the 62 blood samples and on the 125 cfDNA samples). Error rates were estimated at each position of the gene and for each substitution, totaling *1704*3=5,112* values. We were then interested in estimating the contribution of each possible nucleotide change on the error rate. We therefore computed, for each error-rate range *e* in [{10^−5^, 10^−4^};{10^−4^, 10^−3^};{10^−3^, 10^−2^};{10^−2^, 10^−1^}] and for each possible base change *b* in [G>T, C>A,…, A>C, T>G]:

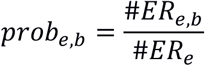

with #*ER_e,b_* being the number of estimated error rates in the class *e* observed for a base change *b*, and *#ER_e_* being the total number of estimated error rates in the class *e*.

In the case of the Ion Torrent sequencing, we observed a sufficiently high number of single nucleotide variations (SNVs) (n=5,112) to also compute the distribution of error rate depending on the 96 possible SNVs taking into account the preceding and following bases to evaluate the effect of the sequence context. Similarly, the high number of insertions (n=7,662) and deletions (n=1,724) detected allowed us to also compute the distribution of estimated error rates (i) as a function of the length of the inserted/deleted sequences; and (ii) as a function of the length of homopolymer regions for the insertion/deletion of one base pair.

### cfDNA and matched tumour analysis for validation

Observed deleterious mutations in the *TP53* gene of a cfDNA lung cancer patient are generally expected to be derived from their tumour (but see Fernandez-Cuesta et al. 2016 Ebiomedicine) (20). Therefore we used the tumour samples as a proxy for validation of the identified cfDNA mutations. To limit our false discovery rate, samples that harbored a high number of raw mutations (>100) in at least one of the two technical replicates were excluded. This removed 4 SCC and 7 SCLC from the 46 matched samples. We considered only cfDNA mutations that passed post-calling filters, *i.e*. a RVSB (Relative Variant Strand Bias) (20) lower than 0.85, no high-VAF variant (*i.e*. a VAF ten times higher than the candidate mutation) within 5 base-pairs upstream or downstream, and a VAF higher than 10% if the mutation is found in a low confidence base change (*i.e*. where technical duplicates don’t cluster together; see Supplementary Methods). We independently performed the needlestack variant calling on the cfDNA samples and the matched tumour samples.

## RESULTS

### Sequencing error rates depend on the alteration type

Globally, 95% of the error rates across alterations were estimated as lower than 10^−2.5^ in both sequencing technologies (Figure 1A). Nevertheless, the error rates varied importantly across the target sequences and alterations. For the amplicon-based Ion Torrent sequencing, transitions had 5-fold higher error rates than transversions (Figure 1A), on average, although not clearly influenced by the sequence context when considering the flanking 3’ and 5’ bases (Supplementary Figure S2). For exome-capture sequencing, a bulk in the distribution of transversion-like errors is observed at an error rate in the order of 10^−2.5^ (Figure 1A). When looking at the proportion of different nucleotide substitutions across multiple ranges of sequencing error rates (Figure 1B), we observed that in this range (10^−2^ – 10^−3^) the majority of substitutions correspond to G>T transversions, previously reported and suggested to be related to DNA sonication(21).

**Figure 1:**
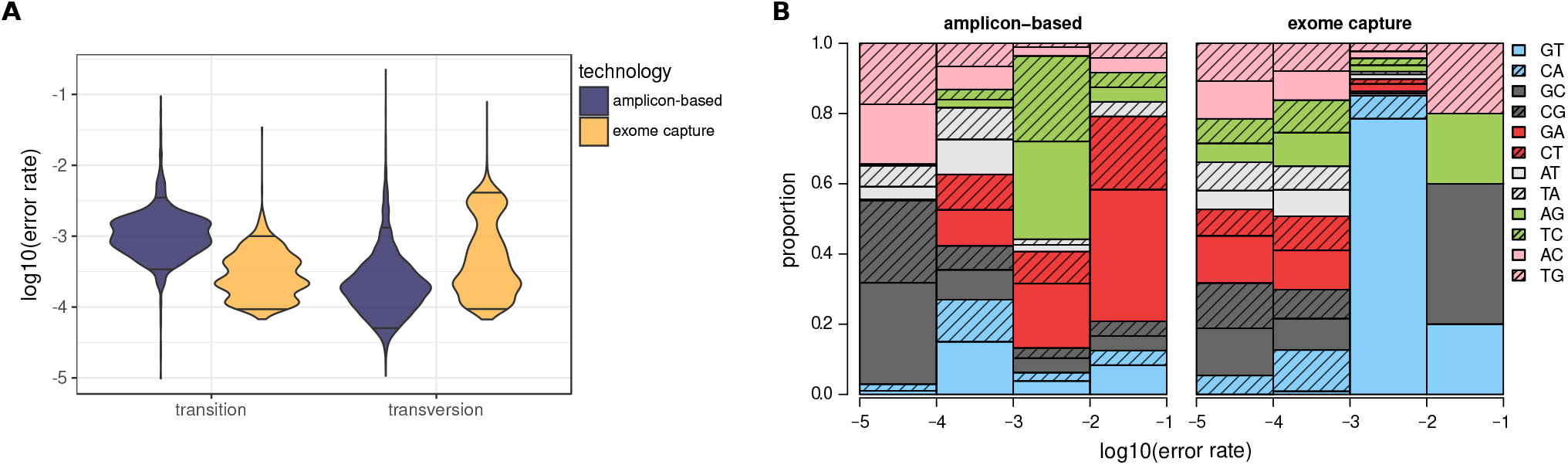
Sequencing error rates estimated by needlestack across the *TP53* gene. (A) Distribution of sequencing error rates in log-10 scale across the 1,704 positions accounting for a total of 5,112 values. Results are stratified by type of base change: transition or transversion (x-axis) and by sequencing technology (IonTorrent Proton amplicon-based data in violet and Illumina exome capture data in yellow). Horizontal black lines correspond to the 5% quantiles of each of the sequencing error rate distribution. (B) Contribution of each of the 12 possible base changes on the estimated error rate. Error rates are stratified by ranges ([10^−5^, 10^−4^];[10^−4^, 10^−3^];[10^−3^, 10^−2^];[10^−2^, 10−^1^], x-axis). Base change contributions are colored according to DNA strand equivalences (*e.g*. G to T and C to A are both colored in blue).

As previously reported, we observed a large number of indels (9,389) in the Ion Torrent sequencing data (22). We found that the error rate is dependent of their length: long indel (with a size greater than 3bp) error rates are around 100-fold lower than 1bp indel error rate (Supplementary Figure S3A). As previously reported (22), the error rate also increases with the length of homopolymer region, reaching 1% for repetitions of four nucleotides (Supplementary Figure S3B).

### Variant detection limit depends on the error rate

Importantly, errors identified in the previous section are classified as such by needlestack, and not as potential variants, even when the error rate is high, as opposed to traditional variant callers which consider samples individually and that rely mostly on the VAF (21). Figure 2A illustrates a position at which needlestack identifies a high error rate (*e*_jk_ = 3.8) without reporting any variant, even though alternate reads are observed in individuals VAF’s up to ~ 9%. Figure 2B illustrates a position with a very different estimated error rate (*e*_jk_ = 10^−4^) where a putative very rare variant is identified. It is also noteworthy that the variant identified in Figure 2B has a VAF ten times lower (10^−3^) than the error rate estimated in Figure 2A, indicating that the sensitivity to detect a variant is considerably improved at the site with the lower error rate, highlighting the need to quantify the error rate distributions for each candidate mutation independently.

**Figure 2:**
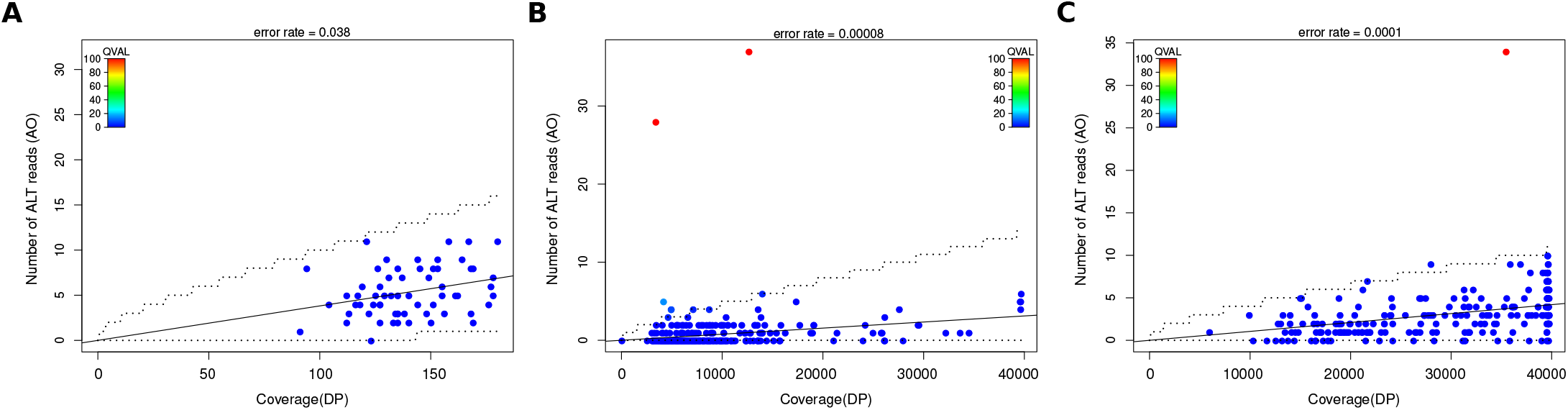
Needlestack regression plot for three independent genomic alterations. Each dot corresponds to a sequencing library of a sample and the dots are colored according to the Q-values attributed by needlestack. Red dots are libraries identified as carrying the mutation by needlestack (their Q-values are higher than 50). Dotted lines correspond to 99% confidence interval around the estimated error rate. (A) Example of a G to T transversion from exome-hydrid capture Illumina sequencing where the sequencing error rate is estimated as 3.8×10^−2^ and no variant is detected. (B) Example of a validated mutation (*i.e*. found in the two technical replicates of the same sample) with a VAF around 10^-3^ with a corresponding sequencing error rate estimated around 10^−4^. (C) Example of a non-validated mutation with a VAF at 10^−4^ in the positive library. Each dot corresponds to the library of a sample and the dots are colored according to the Q-values attributed by needlestack. Red dots are libraries identified as carrying the mutation by needlestack (their Q-values are higher than 50).

### Technical replicates reduce low VAF false calls

We noted that the majority of variants detected by needlestack in the cfDNA of healthy patients harbour a particularly low VAF, typically under 0.5% (Figure 3A, black solid line). Importantly, the majority of these variants are not present in a second library preparation (a technical duplicate) of the same sample (Figure 3A, blue lines). Such variants illustrate an additional type of errors found in NGS data that do not consistently re-occur in the samples and that are not validated when sequencing a technical replicate of the sample, for example those introduced by polymerase chain reaction (PCR) amplification errors. These non-systematic artefacts are not expected to be captured by our error model and should be detected by needlestack as outliers (see Figure 2C for such an example). Importantly, we showed that this high number of calls not validated in a technical replicate of the sample is not dependent on our method (Figure 3A, blue lines). Subsequently, here, for the evaluation of needlestack’s ability to detect efficiently low VAF mutations, we added the condition that variants are also detected in the technical duplicates to account for this type of error (Figure 3A, blue line).

**Figure 3:**
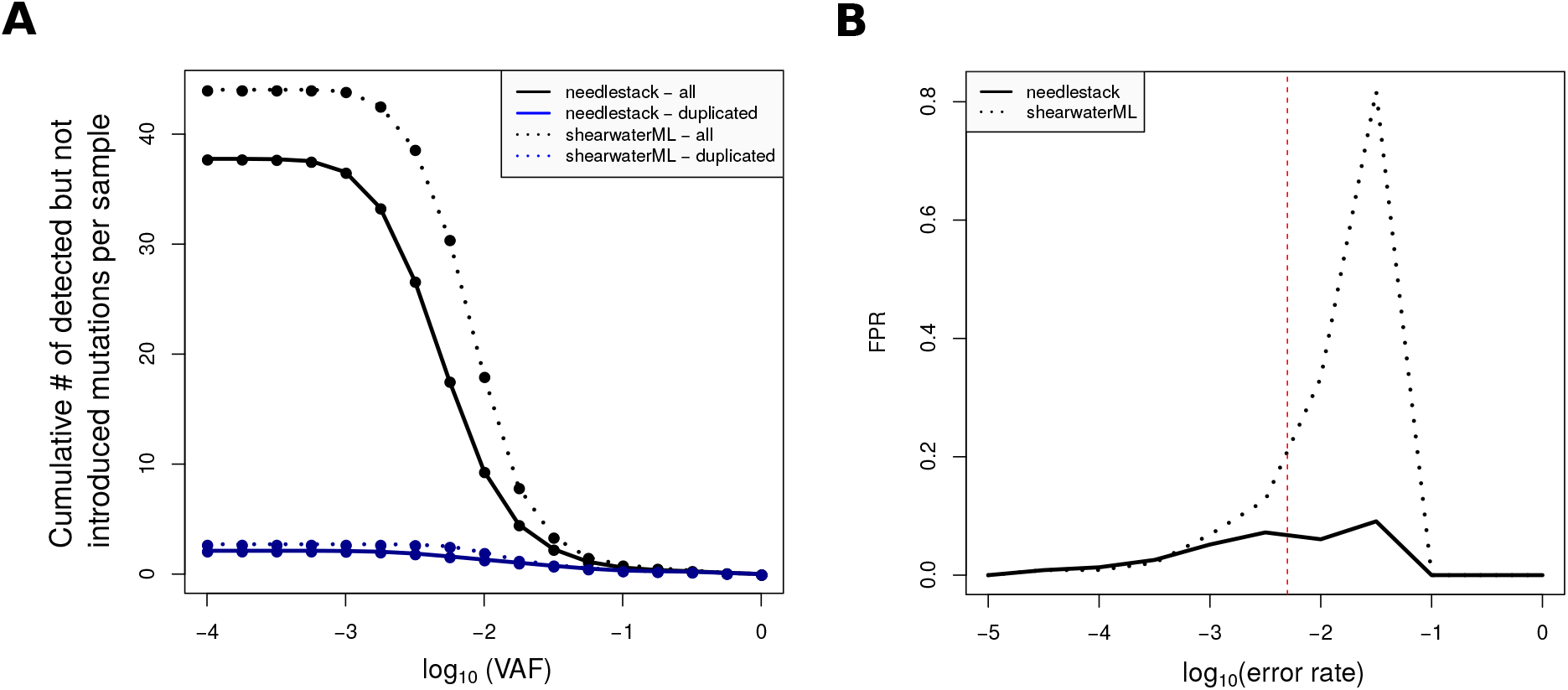
Needlestack and shearwaterML variant calling false discovery overview from *in-silico* simultations with BAMsurgeon on 125 duplicated samples of circulating cell-free DNA from healthy individuals. (A) Cumulative number of detected mutations that were not introduced by BAMsurgeon as a function of the VAF (in log10 scale) of the mutations, for both methods (needlestack in solid lines and shearwaterML in dotted lines). This number is computed as the average per library when considering all mutations (black lines) and as the average per sample when considering validated mutations (*i.e*. found in the two technical replicates of the same sample blue lines). (B) False positive rate (per alteration) for both needlestack and shearwaterML, depending on the estimated error rate at the position (in log10 scale). The red line corresponds to the error rate threshold *t* used for shearwaterML (0.005). ShearwaterML uses this threshold to remove *a-priori* true variants, *i.e*. samples with a VAF>*t*, to then estimate the error rate.

### Performance evaluation using *in-silico* simulation of somatic mutations

From the 10,000 mutations introduced by BAMsurgeon, needlestack detected 5% of mutations with a VAF lower than 0.1%, 51.4% of mutations with a VAF between 0.1% and 1%, and 99.4% of mutation with a VAF higher than 1%. As expected, the sensitivity of needlestack is highly dependent on the sequencing error rate. Indeed, needlestack does not call a mutation if the sequencing error rate for that alteration is greater than or in the same range as the VAF of the candidate mutation (Figure 4). As an example, needlestack detected 0%, 6.5%, and 47.8% of SNVs with a VAF of 0.1% at positions where the sequencing error rate was higher than 0.1%, between 0.1% and 0.01%, and lower than 0.01%, respectively. When comparing needlestack and shearwaterML, we found that globally needlestack sensitivity was higher than that of ShearwaterML, and, for example, ShearwaterML detected 7.7% of all inserted mutations with a VAF at 10^−3^ whereas needlestack detected 16.8% of these mutations. Given *t* the shearwaterML *a-priori* threshold on the sequencing error rate (Figure 3B red line) and *e* the observed sequencing error rate, we showed that the false positive rate of shearwater is markedly increased when *t>e*, whereas needlestack’s false positive rate is stable across the whole range of error rates (Figure 3B).

**Figure 4:**
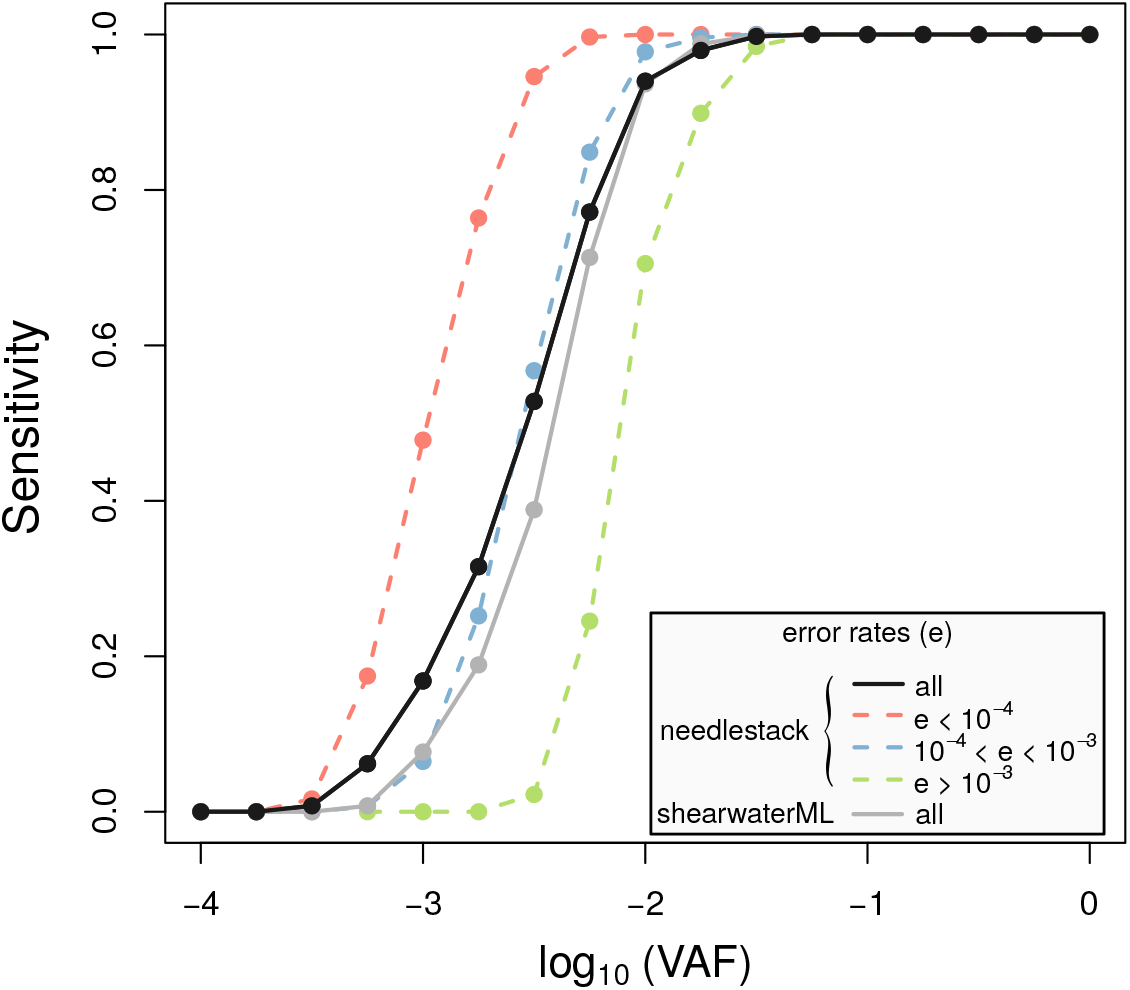
Performance of needlestack for somatic mutation calling using simulated data. The sensitivity of needlestack is shown for multiple values of VAF (in log10 scale, x-axis) of *in-silico* simulated mutations. A total of 10×1,000 SNVs were introduced using the BAMsurgeon software, on a set of 125 samples sequenced at the *TP53* gene with the IonTorrent Proton technology. Needlestack sensitivity was computed independently for different error rate ranges (e, red, blue and green lines). Black line corresponds to the global sensitivity for all the mutations independently of the sequencing error rate. Global sensitivity of shearwaterML for the same data is shown in grey.

### Detection of tumour-derived mutations from cell-free DNA

Next, we tested needlestack’s ability to detect very low VAF mutations in a biologically relevant setting. For this we screened cfDNA extracted from plasma samples from 35 lung cancer patients where the matched tumour sample was analyzed concurrently, and considered the concordance between the identified variants. A total of 22 *TP53* mutations from 18 samples (9 SCLC and 9 SCC) where identified in the cfDNA. 16/22 (70%) mutations were called in the tumour of the same patient. All the 12/22 cfDNA mutations considered as deleterious (*i.e* indels, non synonymous SNVs with a REVEL score higher than 0.5, stopgain or stoploss variants) (23) were present in the tumour. cfDNA and tumour VAF were found to be moderately correlated, which is concordant with previously reported results (24) (Pearson correlation coefficient ρ equals to 0.59, Supplementary Figure S4A). Details of the 22 cfDNA mutations and corresponding observations in the tumour matched samples are provided in Supplementary Table S1. The needlestack plots of a low VAF cfDNA mutation validated in the tumour are shown in Supplementary Figure S4B.

### Application to germline variant calling

For rare germline variants from 33 whole exomes, needlestack has a sensitivity of 95.64% to detect non-reference genotypes when using bead array data as a gold standard, which is quite similar to the GATK-HC Haplotype Caller results (95.48%). GATK-HC and needlestack variants concordant with the bead array (19,515 of the 20,439 variants) had VAF distributed around 50% and 100%, as expected for germline variants (Figure 5A). Most of the few calls that were not validated in the array were also centred around 50% and found by both variant callers, implying that they certainly contain additional heterozygotes that the SNP array failed to detect. Finally, the majority of variants not identified with NGS had no sequencing reads supporting the alternative allele detected by the array (841/892 variants), suggesting that these variants are potentially false positive results from the SNP array (Figure 5A).

**Figure 5:**
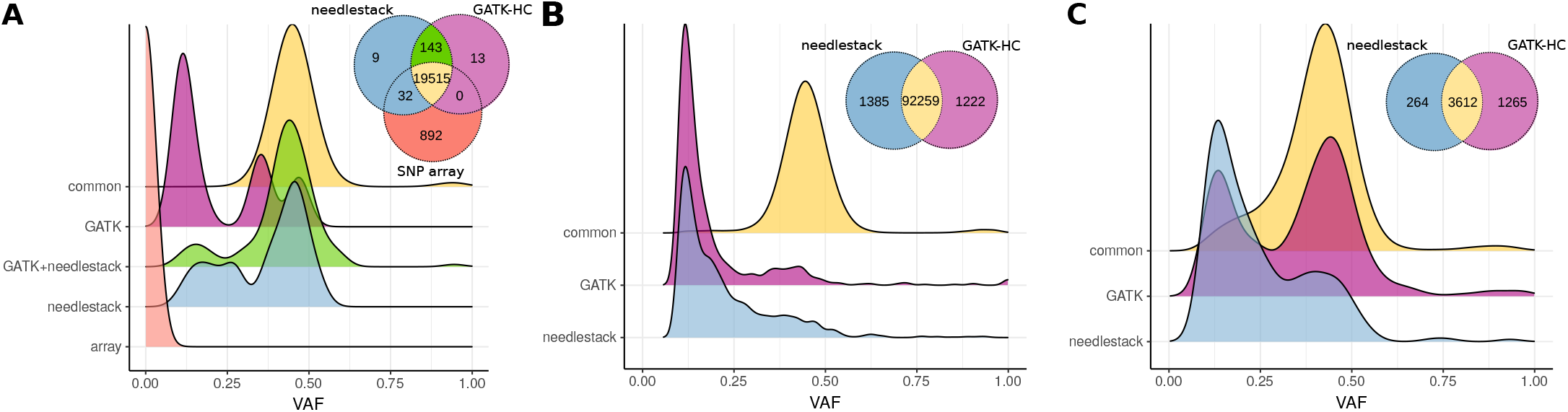
Germline variant calling comparison between needlestack and GATK-HC across 62 samples. Both distributions of the VAF and Venn diagrams showing the concordance of called mutations are shown. VAF distributions are colored according to the Venn diagram. (A) Comparison between both methods and an Illumina bead array containing gold standard genotypes available for a total of 33 samples. (B) and (C) Comparison between needlestack and GATK-HC called mutations without any reference gold standard for both SNVs (B) and indels (C).

Because SNP arrays are biased toward sites amenable to the design of Illumina BeadArrays (25), we also undertook needlestack and GATK germline genotyping of SNVs and indels calls across 62 exomes. Respectively 97.3% and 70.3% of the SNV and indel calls were concordant (Figure 5B-C) with VAFs around 50%, whereas the genotypes identified uniquely by one of the two methods tended to have low VAF. For indel calling, 46% of calls unique to needlestack and 34% of calls unique to GATK are more than 10bp long, compared to only 12% of common calls. This suggests that discrepancies among the methods can be partially explained by longer indels that are difficult to align and call. For 66% of uniquely called indels by GATK-HC, no alternate reads were present in the BAM file used by needlestack, suggesting divergences in the assembly steps (haplotyper Caller versus ABRA). Interestingly, for 52% of the SNVs detected by GATK-HC and not by needlestack, needlestack estimated an error rate higher than 1%, pointing to possible false positives in the GATK calls (Supplementary Figure S6).

## DISCUSSION

The needlestack method is based on the notion that, as error rates strongly vary along the genome, their dynamic estimation from multiple samples, for each potential base change at a given DNA position, may assist in accurately identifying sequence variants. Here, we have demonstrated that, even within a single gene (*TP53*), and even if the sequencing error rate is generally low, it varies importantly across positions and base changes (Figure 1). Needlestack implements a robust negative binomial regression for this purpose, and the ability of the method to identify variants will be dependent upon the error rate at that particular site and for that base change. By identifying sequence variants as outliers relative to the error model, needlestack maximizes the sensitivity to detect variants in a dynamic manner relative to the error rate in that particular setting. As such, low allelic fraction variants are identified from sites with low errors rates, whereas in settings where error rates are high, needlestack maintains reasonable false discovery rates (Figures 3 and 4).

We have benchmarked our method using both simulated and real data from different sequencing platforms. First, we have tested our method on low VAF mutations using BAMsurgeon to generate *in-silico* mutations and have compared our findings to variants identified by a similar rare variant orientated algorithm shearwaterML (9,10). We have shown that our method outperforms shearwaterML for VAF lower than 10^−2^ and that the performance of shearwaterML highly depends on the difference between the error rate *e* and the error rate *a-priori* threshold *t* (see methods for details). Contrary to shearwaterML, needlestack’s false discovery rate is not dependent on the sequencing error rate. In addition, needlestack also considers indel mutations. For this type of variant, the sensitivity of needlestack is slightly reduced compared to SNVs (Supplementary Figure S5A-B). This is potentially due to the increased complexity of the assembly step around indels compared to SNVs. Moreover, needlestack detects a high number of indels replicated in the two technical duplicates that were not *in-silico* introduced (around 8 by samples in average), whereas *TP53* is not expected to harbour many indels in healthy patients. These mutations can be moderated using a filter on the strand bias, as previously reported (Supplementary Figure S5C-D) (26).

The true specificity of needlestack cannot be achieved with BAMsurgeon simulations, due to a probably very low presence of true mutations in the cfDNA of healthy patients that is difficult to determine *a priori* (20). We therefore have estimated the validation rate in the tumour of deleterious cfDNA mutations identified by needlestack in 35 lung patient cfDNA samples. All of these 12 mutations were validated in the tumour.

Finally, we have benchmarked needlestack on germline mutations using SNP array data to validate the mutations detected in WES of 33 individuals, and showed an excellent concordance when results are compared with both a SNP array as a gold standard set and calls from GATK HaplotypeCaller (19). This illustrates that needlestack, even if based on a totally different approach to detect variants, can reach similar performance to state-of-the-art germline variant callers.

The needlestack method nevertheless has several limitations. Even though needlestack is extremely sensitive, it is suited to detect rare mutations rather than common germline variants or highly re-occurring hotspot mutations. Adding an *a-priori* threshold for the error rate (extra_robust mode – see Supplementary Methods) can partially offset this limitation, but is only applicable to particular situations for the reasons explained above. More importantly, the inherent logic of the needlestack approach corrects for errors that have a tendency to reoccur, as such errors that are rarer are identified as outliers in the regression. Following this, needlestack does not correct for sample-specific artifacts such as (i) (sample specific) stochastic alignment errors and we recommend to use it in conjunction with an assembly based re-alignment method (27); (ii) polymerase errors introduced in PCR amplification step; (iii) complex errors leading to features like strand bias. Such errors remain a feature in NGS data (Figures 2C and 3A), thus additional error correction (28,29) and/or validation techniques are needed. This can be achieved with hard filtering on the output statistics such as the VAF or the strand bias, but also with machine-learning-based approaches applied to multiple variant summary statistics when validated data are available to inform the model. Here we have controlled for these errors by undertaking technical duplicate of each sample and conditioning on the requirement that the variant must be present in each preparation.

Our pipeline is implemented using nextflow (12), to facilitate its scientific reproducibility but also efficient parallel computations. Needlestack is also provided with Docker (30) and singularity (31) containers to avoid installation of dependencies and produce perfectly reproducible results. Needlestack is a user-friendly pipeline that can be run in one command line. In addition, needlestack implements a power calculation to estimate if the coverage is sufficient to call a mutation (see Supplementary Methods for details). Using this power analysis, it can predict the germline or somatic status of a mutation when applied to tumour-matched normal mode. This also allows needlestack to flag mutations with an “unpredictable” status (when the coverage is to low) to accurately control the false discovery rate. Source code is available on GitHub and is versioned using a stable git branching model. Importantly, this approach is relatively computationally efficient and parallelisable. This allows error models to be built even across large target stretches of DNA, enabling applications at the exome level, genome levels or to most forms of sequencing data. As an example, needlestack takes around 20 hours to analyze 100 WES when launched on 100 CPUs.

In summary, needlestack uses a robust model of sequencing errors to accurately identify DNA mutations potentially in very low abundance. The model takes the advantage of batch sequencing of multiple samples to precisely estimate the error rate for each candidate alteration. Needlestack can be applicable to various types of studies such as cfDNA, histological normal tissue investigation or high precision tumour subclonality estimation by providing a high sensitivity for low allelic fraction mutations.

## AVAILABILITY

needlestack is an open source software and is available in the GitHub repository (https://github.com/IARCbioinfo/needlestack).

## ACKNOWLEDGEMENT

We would like to acknowledge Alain Viari, Dariush Nasrollahzadeh Nesheli and David Muller for their helpful inputs and feedbacks.

## DISCLAIMER

Where authors are identified as personnel of the International Agency for Research on Cancer/World Health Organization, the authors alone are responsible for the views expressed in this article and they do not necessarily represent the decisions, policy or views of the International Agency for Research on Cancer/World Health Organization.

## SUPPLEMENTARY MATERIAL

### SUPPLEMENTARY METHODS

#### Robust Negative Binomial regression

The original method (1) was established on falls data where the predictor variable took values from 0 to a couple of hundreds. Here we need to take into account cases where we sequenced deeply and therefore the predictor variable DP can be up to hundreds of thousands. The first model uses integrals of bounding functions for the maximum likelihood estimation (MLE) of *e*_jk_ to keep robustness, which can take a very long time for high coverage data. To save computing time, we approximate the calculation of integrals required for the MLE of *e*_jk_. Instead of computing the sum of all values corresponding to the integral, we interpolate the points using the *spline* function in R, and compute the sum of a set of sampling points with a reasonable size (default is 100).

The initial estimation of *e*_jk_ for the MLE algorithm is based on a Poisson model, and because of a lack of robustness the following MLE of *σ*_jk_ can take a lot of time. We thus define our initiation of *e*_jk_ as the mean of observed *e*_ijk_ after passing the Tukey’s outlier filter, *i.e* an observed *e*_ijk_ is taken into account for the mean computation if and only if it verifies *e*_ijk_ ≤ *Q*_3_ + 1.5 * IQR, with Qi=i^th^quartile and IQR=Q_3_ − *Q*_1_.

#### Implementation

Needlestack is implemented as one major process, which can be executed in parallel for multiple input chunks, each corresponding to a set of genomic positions. This process is defined as a chain of piped commands: firstly, it runs samtools mpileup utility to compute, for each of the input BAM files, the list of read nucleotides overlapping the input positions. Then, it translates the samtools output into an easier to process format through a custom C++ tool (mpileup2readcount). Finally, needlestack uses its own R script to run the variant calling independently at each position and for each observed alternative base change, and produces a resulting VCF file that will be merged with other created VCF if run in parallel mode. See Supplementary Figure 5 for details on the pipeline. Needlestack is written in the nextflow (2) domain-specific language, allowing high scalability and reproducibility, but also efficient parallel execution. Needlestack source code is freely available on Github (3), and a Docker container image is hosted on DockerHub (4). This docker image is based on conda and Bioconda (5), a sustainable and comprehensive collection of bioinformatics software that help to easily install workflow dependencies.

#### Tumour-Normal pairs method

We have implemented, in addition to our basic model, a method to classify any observed variant as somatic or germline when needlestack is launched in tumour-normal pairs mode (Supplementary Table S2). For this, for each variant detected only in either the normal or tumor sample, needlestack estimates the power to detect it in the other sample. Indeed, not observing a variant in the other sample could be due to the lack of power to detect it, in particular when the depth of coverage is not sufficient. Our power metric is based on the expected Q-value of the variant if truly present, which depends on the observed coverage of the sample at the site, the expected allelic fraction and the error model estimated by Needlestack. If this Q-value is below the user-chosen threshold, we report a lack of power. For a particular individual sequenced for a tumour-normal pair, needlestack classifies its variants as follows: if a variant is observed in the tumour sample, it is classified as “somatic” if not observed in the normal although the power was sufficient, and as “unknown” in a case of a lack of power in the normal sample. If a variant is observed in the normal sample, it will be labeled as “germline”, and sub-labeled as “confirmed” if also found in the tumour, “unconfirmed” if not found in the tumour whereas power was satisfactory, and “unconfirmable” if there was not enough power in the tumour to detect it if present.

To obtain the expected Q-value of a variant, the first step is the computation of the minimum expected alternative read count (6) in the other sample (tumor or normal). The expected AO of a (germline) variant in the normal sample is computed as follows:

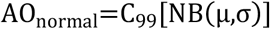

with *C_99_* the 99^th^ centile, *NB*=negative binomial distribution, *μ*=0.5 x coverage at the position, and *σ*=dispersion parameter (by default=0.1).

The expected AO of a variant in the tumor sample is computed as follows:

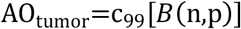

with *C_99_* the 99^th^ centile, *B*=binomial distribution, *n*=coverage at the position, and *p*=minimum variant allelic fraction expected (by default=0.01).

Then, given this expected AO, we compute the expected p-value that corresponds to the probability to belong to the error model that needlestack has estimated. Finally, we transform this p-value into a Phred-scale Q-value to obtain the expected Q-value of the variant.

#### cfDNA and tumour sequencing for *in-silico* simulations and tumour validation

CfDNA was extracted from 0.8-1.3 mL of plasma using the QIAamp DNA Circulating Nucleic Acid kit (Qiagen) following manufacturer’s instructions. CfDNA was eluted into 100 μL of elution buffer and quantified with the Qubit DNA high-sensitivity assay kit (Invitrogen Corporation). Twenty-one amplicons of 150 bp in size were designed (Eurofins Genomics Ebersberg, Germany) to cover exons 2 to 11 of *TP53*. The GeneRead DNAseq Panel PCR Kit V2 (Qiagen) was used for target enrichment. A validated in-house protocol was used to set up multiplex PCRs in 10 μL reaction volume, containing 5 ng cfDNA, 60 nM of primer pool and 0.73 μL of HotStarTaq enzyme. The experiments were carried out in two physically isolated laboratory spaces: one for sample preparation and another one for postamplification steps. Amplification was carried out in a 96-well format plates DNA engine Tetrad 2 Peltier Thermal Cycler (BIORAD) as follows: 15 min at 95°C and 30 cycles of 15 seconds at 95°C and 2 min at 60°C and 10 min at 72°C. Two technical duplicates were undertaken for each cfDNA sample including amplification, library preparation, and sequencing. Each technical duplicate pair was assessed on two separate plates to limit the possibility of a contamination.

For the tumour sequencing, eighty nanograms of each DNA sample was used as template to set up four separate PCR reactions (20ng/pool) using the Qiagen GeneRead DNAseq Panel PCR Kit V1 and primer mix (Qiagen), following manufacturer’s instructions. The amplified PCR products were then pooled, purified with the Serapure magnetic beads and subjected to library preparation including adapter ligation, purification, and amplification using the NEBNext Fast DNA Library Preparation Kit (New England Biolabs). About 200 ng of individual libraries were pooled into a single tube and size selection (230~250 bp) of pooled libraries was performed using 100μL aliquot of pooled libraries onto a 2% agarose gel and MinElute Gel Extraction Kit (Qiagen).

Template preparation was done on the Ion OneTouch2 instrument using the Ion PGM Template OT2 200 Kit, followed by sequencing on an Ion Torrent PGM sequencer using the Ion PGM Sequencing 200 Kit v2 (Life Technologies), aiming for mean depth of 500X.

#### Bioinformatics processing

Short reads from NGS sequencing were aligned to the hg19 human reference genome using the Torrent Suite software (v4.4.2) with default parameters. Somatic mutations were detected with needlestack using the version 1.0 and a QVAL threshold at 50. As recommended by Martincorena *et al*. (7), we used a threshold of 20 for the shearwaterML statistic.

Only cfDNA mutations detected in a high confidence base change were considered. A high confidence base change satisfied *P*<0.05 with *P* the probability that the number of duplicated mutations under the null hypothesis would be greater than or equal to the observed number of duplicated mutations, and is given by:

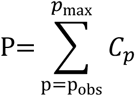

with *C_p_* corresponding to the probability of observing *p* pairs when randomly picking *k* elements from a total of *2N* paired elements calculated as:

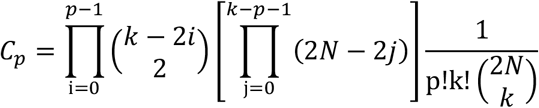

with *N* the number of sequenced samples in duplicates (2N sequenced libraries in total), *k* the number of libraries with a mutation called by needlestack, *p_obs_* the number of duplicated called mutations, and *p_max_* the total number of possible pairs when picking up *k* elements from a total of *N* pairs of elements. The detailed source code for cfDNA mutation analysis including all quality filtering step description is available on GitHub at: https://github.com/IARCbioinfo/target-seq.

For the germline analysis, GATK-HC variant calling was performed using version 3.4 and the HaplotypeCaller algorithm, followed by the joint genotyping step (8). Finally, Variant Quality Score Recalibration following the GATK best practices was applied, using dbSNP 138, HapMap 3.3, 1000 Genomes phase 1 and OMNI 2.5 databases. Options provided were “-tranche 100 -tranche 99.9 -tranche 99.0 -tranche 90.0” for both INDEL and SNP modes. GATK-HC variant calls were filtered on PASS and on Phred-scaled likelihood (8) larger than 20 and on VAF>0.1. BAM files were locally reassembled with ABRA version 1.0 (9) before launching the variant calling by needlestack. Needlestack germline calling was launched using our default germline parameters, *i.e*. QVAL>20, VAF>0.1 and the option –-*extra-robust*. This option helps needlestack to correctly estimate the error rate by excluding common germline variants (defined when more than 10% of the samples with a VAF higher than 20%) that tend to bias this estimation towards high values. For each of these base changes independently, this process first eliminates these germline samples and then estimates the error rate on remaining samples. In this germline analysis, both positions and variants with respectively a median coverage and an individual coverage less than 50 were removed from the whole analysis. Coverages were computed with samtools mpileup, counting only reads with a mapping quality higher than 20 and a base quality higher than 13. We considered as variant frequency the maximum proportion of samples carrying the variant estimated by both methods and then filtered out germline variants with a frequency higher than 10% to consider only rare variations.

#### Computation of ShearwaterML statistic used in the BAMsurgeon *in-silico* simulations

We launched the shearwaterML algorithm on the simulation data sets to compare its global performance with needlestack. As recommended by Martincorena *et al*. (7), we used a threshold of 20 for the shearwaterML statistic. We used default thresholds except that we increased *maxvaf* to 1 to call all mutations, and set *truncate* to 0.005 to avoid true mutations present initially at low VAF to enter the background error model and potentially reduce the sensitivity, as recommended in Martincorena *et al*. (7). ShearwaterML produced *p*-values instead of the shearwater Bayes factor, that we corrected for multiple testing using the Benjamini-Hochberg method which produces then Q-values that we finally transformed into Phred scale *Q*-values (QVAL).

#### Using needlestack to compute the error rate distribution

Needlestack can be forced to compute the error rate for the three possible single nucleotide changes at every query position (i.e. to output the results of the error model even when no variants are identified). This can be achieved by launching needlestack with parameters --all_SNVs, --min_ao 0 and --min_dp 1. This way, in our analysis, the 5112 error rates across the *TP53* gene were computed.

## SUPPLEMENTARY TABLE AND FIGURE LEGENDS

**Supplementary Figure 1:**
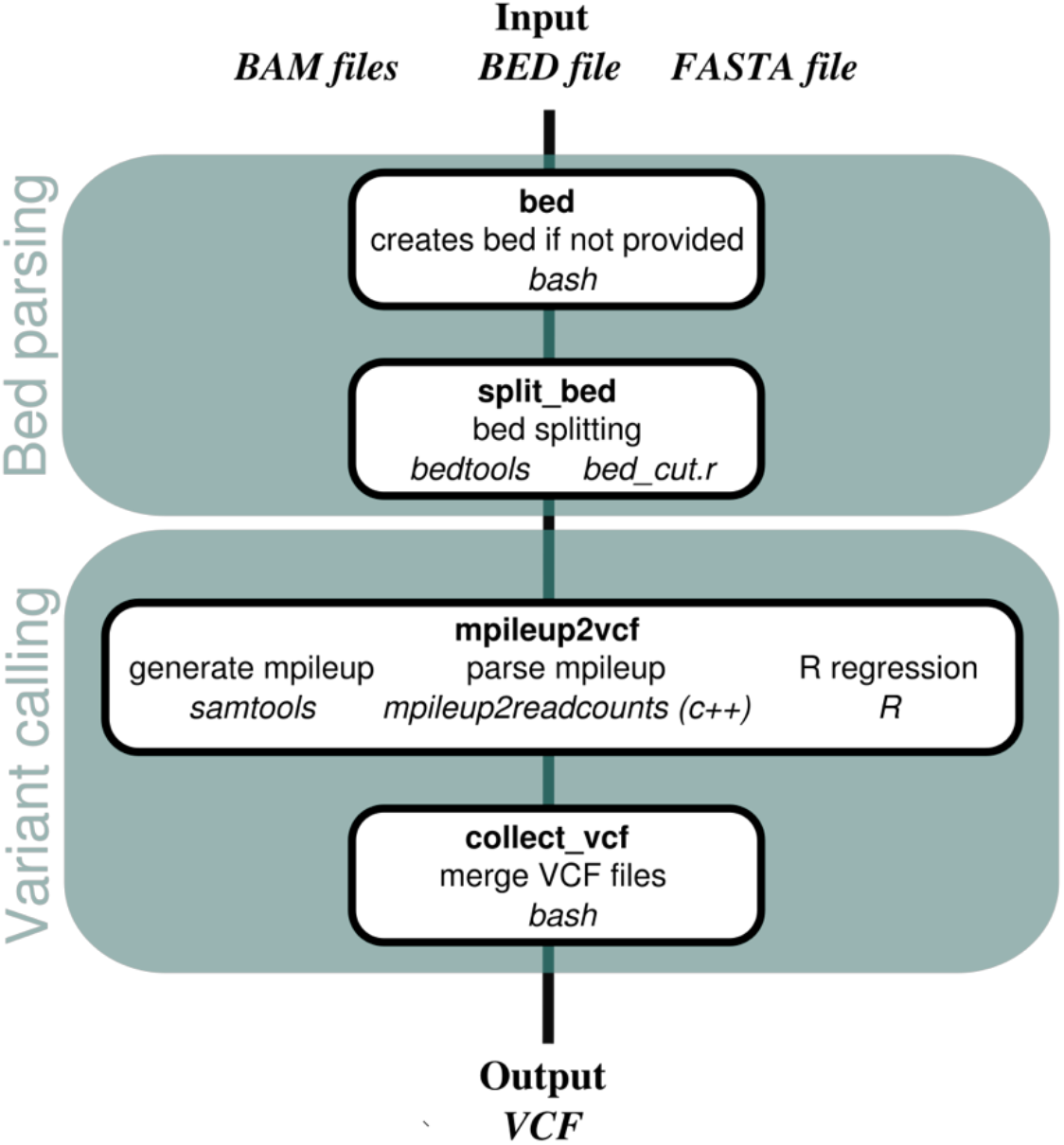
needlestack workflow description. The first step corresponds to the creation of a BED file containing the DNA positions on which the calling should be launched using the fasta index of the input reference genome; this step is optional, only performed if target positions are not provided by the user. The second step splits the BED file into multiple sets of positions to run the algorithm independently on each set in parallel; the number of position sets is given as an input by the user. The third step runs the variant calling from three piped substeps: (i) the mpileup file building using samtools, (ii) the parsing of the mpileup to produce count data per sample in a tabulated readable file, (iii) and the robust regression in R on each tested mutation. The fourth and last step merges VCF files previously produced in parallel and outputs the global result. Workflow orchestration is done thanks to the nextflow domain specific language.

**Supplementary Figure 2:**
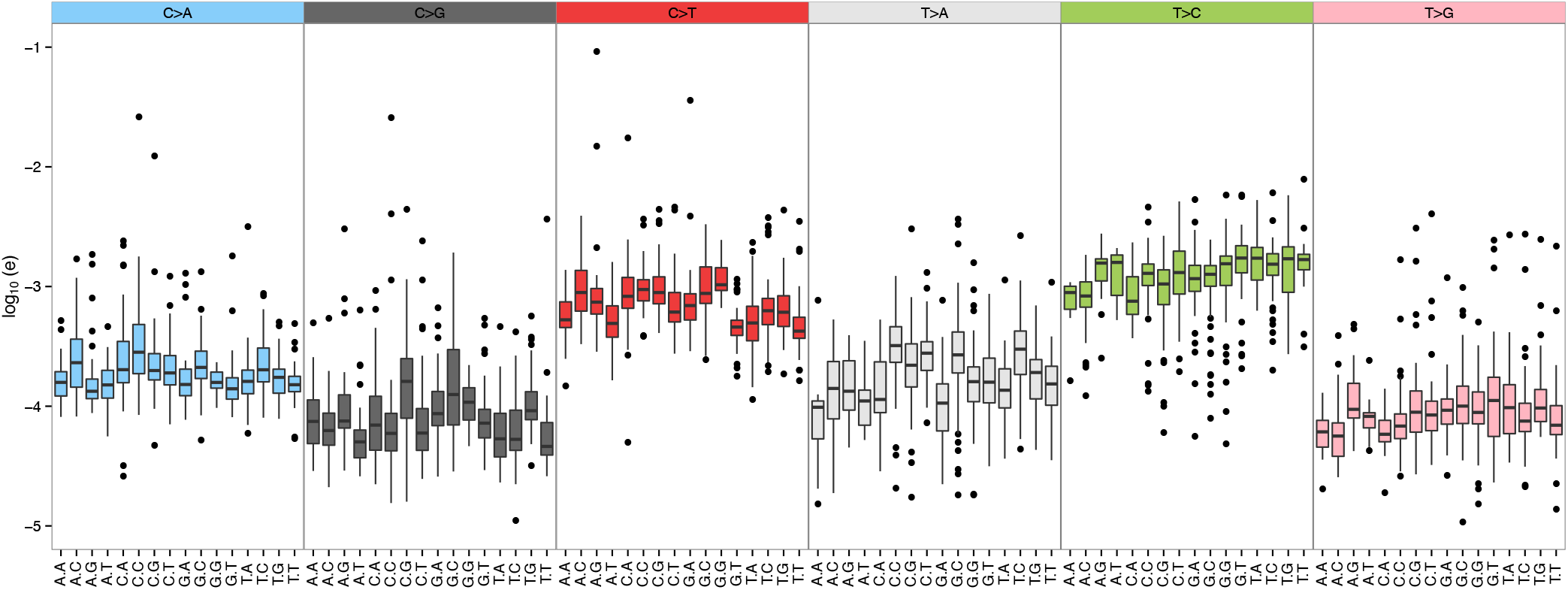
estimated error rate distributions from amplicon-based sequencing of *TP53* gene (median coverage around 10,000X). Distributions of error rates are shown for each of the 96 possible base variation (with flanking 3’ and 5’ bases), and are colored by DNA base changes.

**Supplementary Figure 3:**
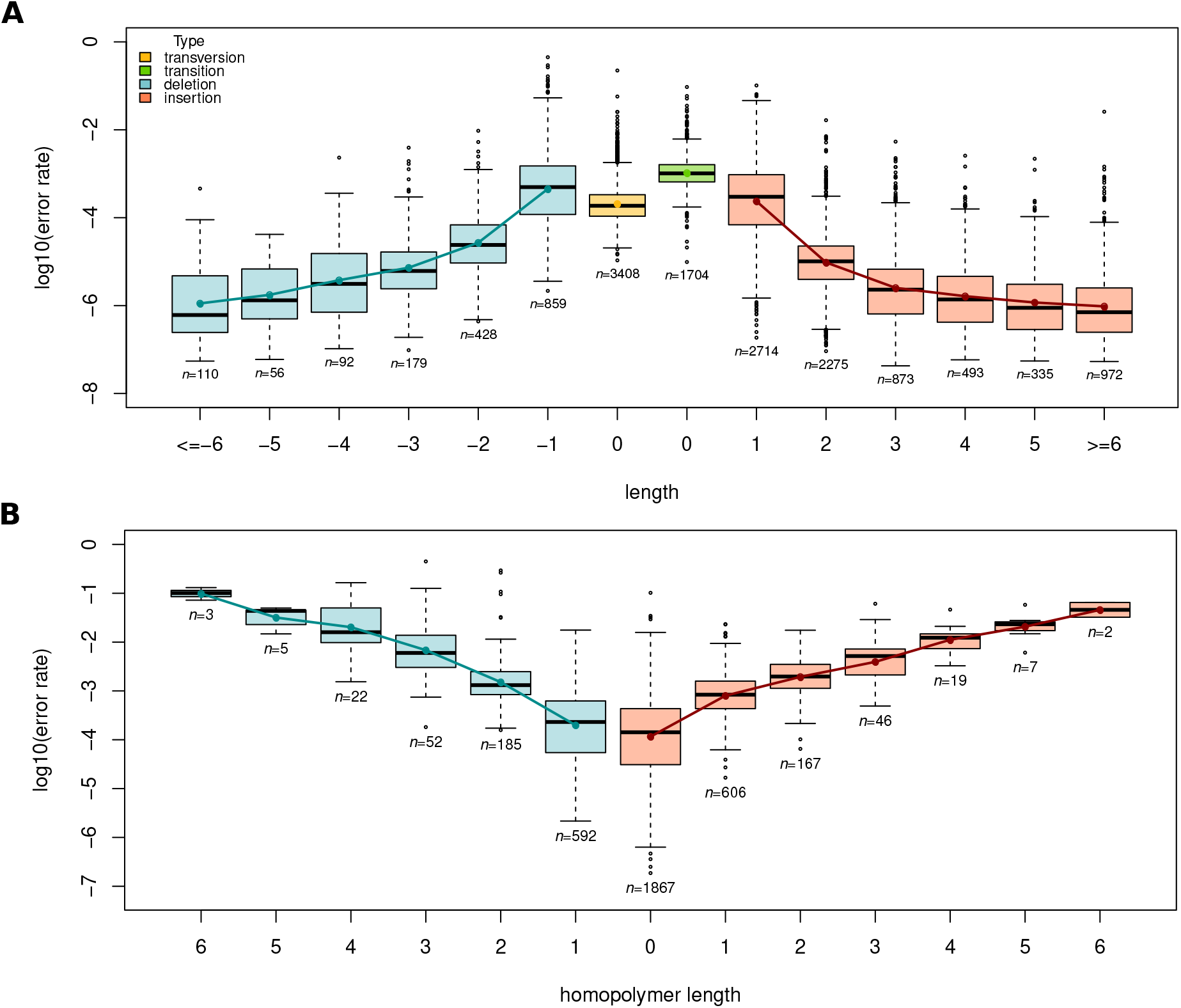
estimated error rate distributions for both SNVs and indels from the same data as used in supplementary Figure 2. (A) Error rate distributions are shown as a function of the type of SNV (yellow for transversions and green for transitions), the length of the insertion (pink) and the length of the deletion (blue), with *n* indicating the total number of error rates used to compute the distribution. (B) Error rate distributions restricted to insertions and deletions, as a function of the size of the homopolymer region (i.e. depending on the number of repeated nucleotides) at the position.

**Supplementary Figure 4:**
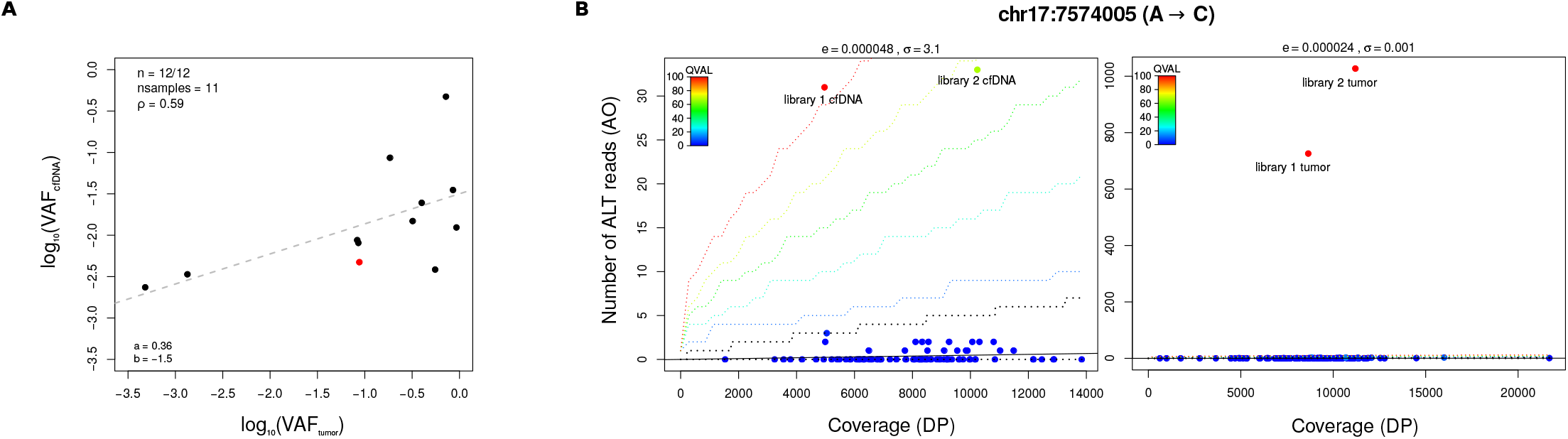
validation of cfDNA mutations in the matched tumour sample for a total of 11 SCLC cases and 24 SCC cases. (A) Correlation of cfDNA and tumour VAF for deleterious validated mutations (total=12). Pearson correlation coefficient ρ was estimated as 0.59. Grey dashed line corresponds to the fitted linear regression characterized by the values *a* (slope) and *b* (intercept). (B) Needlestack regression plots of a validated deleterious SNV (red dot in A). Left panel corresponds to cfDNA data and right panel to tumour data.

**Supplementary Figure 5:**
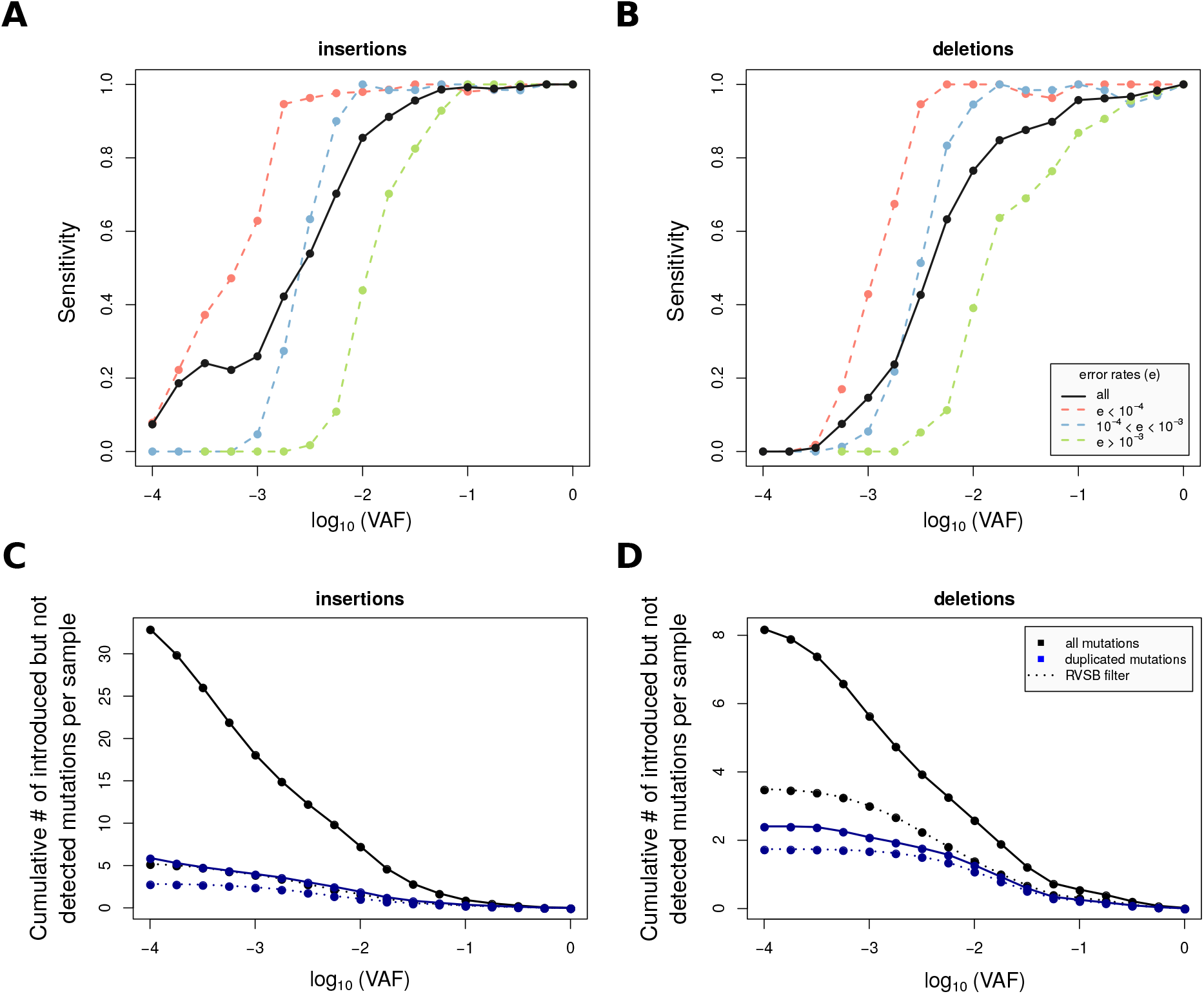
sensitivity of needlestack as a function of the VAF for the *in-silico* simulated insertions (A) and deletions (B), depending on the error rate for the alteration. Cumulative number of false discoveries per sample is shown for insertions (C) and deletions (D) per sample, depending on the VAF of the detected mutation. This number was computed firstly for all detected indels (the result is per library) and secondly for indels validated in the second library (the result is per sample). Dashed lines correspond to the number of indels not introduced by BAMsurgeon that are however not in strand bias (RVSB<0.85).

**Supplementary Figure 6:**
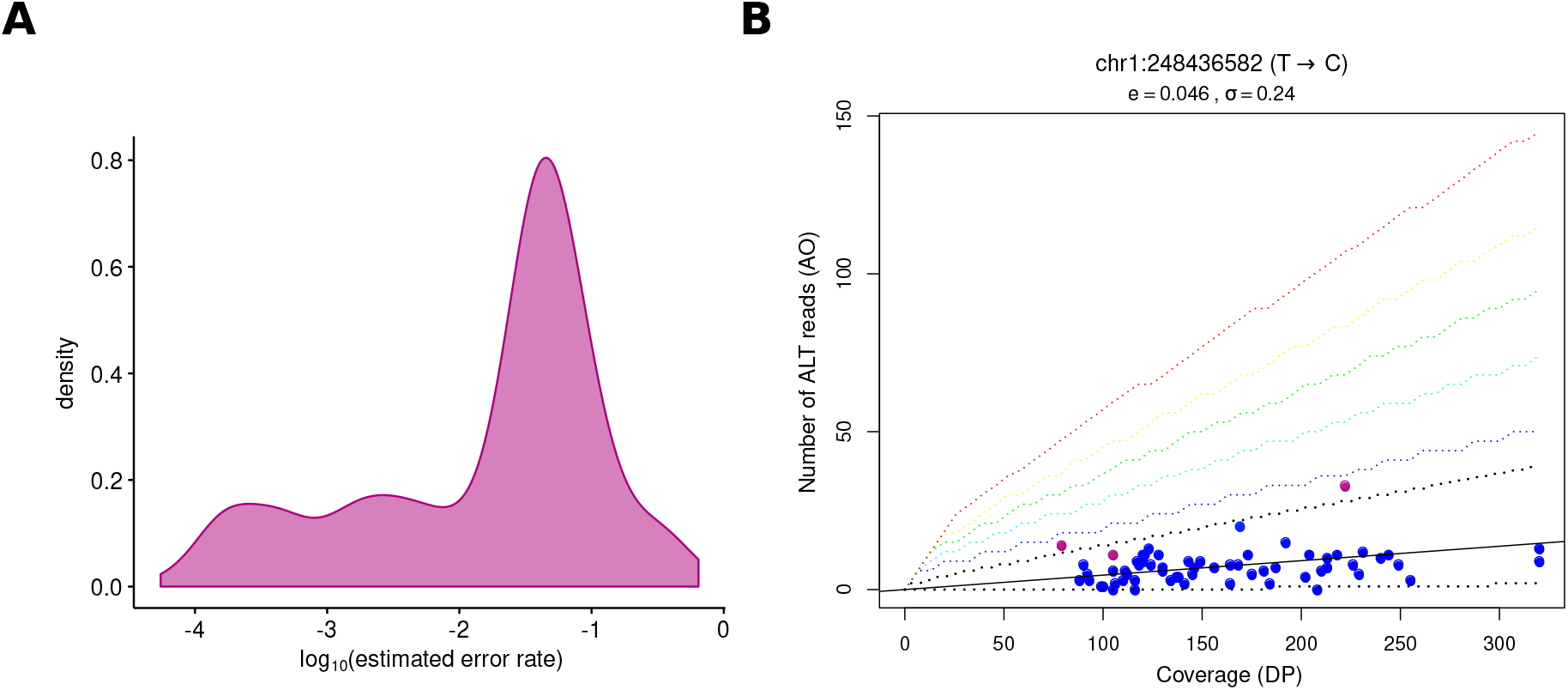
(A) Distribution of the sequencing error rates for SNVs detected by GATK-HC but not detected by needlestack from 62 WES samples (total of 1385 mutations), estimated using kernel density estimation. (B) Needlestack regression plots for one particular position where GATK called 3 variants (in purple). Needlestack estimated a high sequencing error rate (around 1%) for this mutation and therefore did not call it, highlighting the fact that estimating the systematic error rate across multiple sample should reduce the false discovery rate of the method.

Supplementary Table S1: description of the 22 mutations identified by needlestack in the cfDNA of 35 lung cancer patients.

**Supplementary Table S2:**
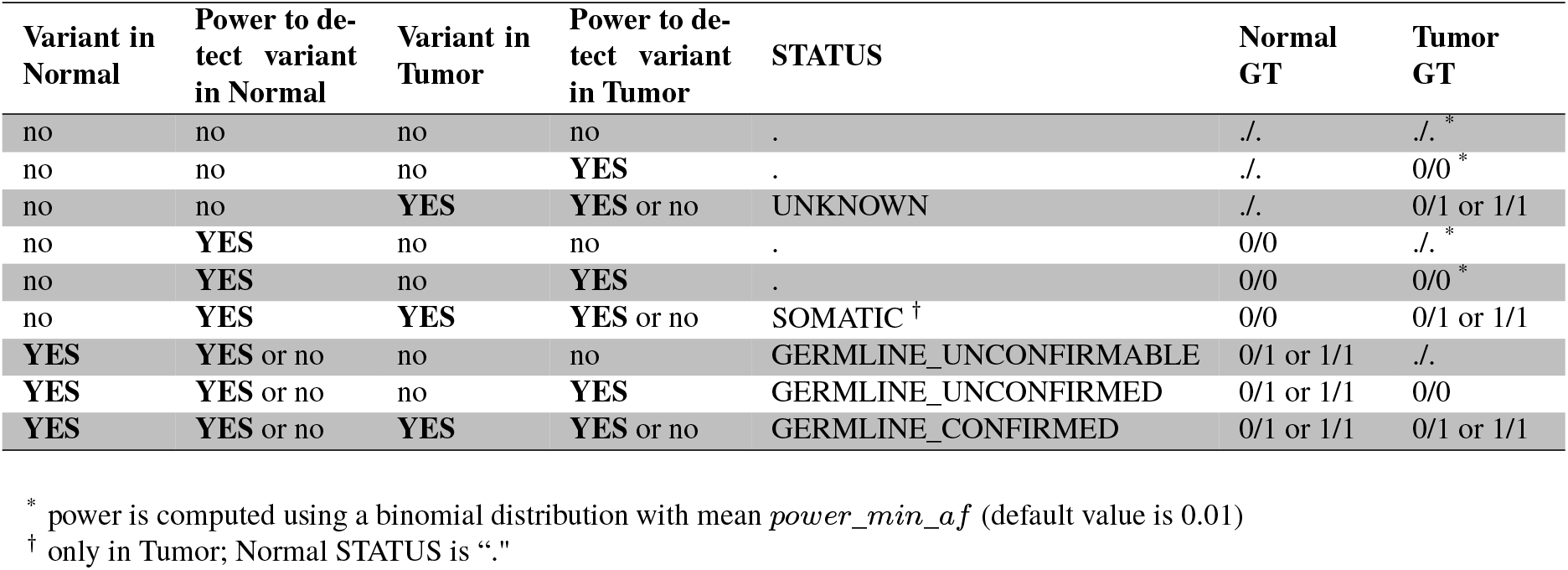
variant status and genotype attributed by needlestack as a function of variant detection and the power to detect variants in tumour and matched normal samples.

| Variant in Normal | Power to detect variant in Normal | Variant in Tumor | Power to detect variant in Tumor | STATUS | Normal GT | Tumor GT |
| --- | --- | --- | --- | --- | --- | --- |
| no | no | no | no | . | ./. | ./.* |
| no | no | no | YES | . | ./. | 0/0* |
| no | no | YES | YES or no | UNKNOWN | ./. | 0/1 or 1/1 |
| no | YES | no | no | . | 0/0 | ./.* |
| no | YES | no | YES | . | 0/0 | 0/0* |
| no | YES | YES | YES or no | SOMATIC<sup>†</sup> | 0/0 | 0/1 or 1/1 |
| YES | YES or no | no | no | GERMLINE_UNCONFIRMABLE | 0/1 or 1/1 | ./. |
| YES | YES or no | no | YES | GERMLINE_UNCONFIRMED | 0/1 or 1/1 | 0/0 |
| YES | YES or no | YES | YES or no | GERMLINE_CONFIRMED | 0/1 or 1/1 | 0/1 or 1/1 |

\* power is computed using a binomial distribution with mean *power\_min\_af* (default value is 0.01)
<sup>†</sup> only in Tumor; Normal STATUS is “.”

## REFERENCES

1. Alioto, T.S., Buchhalter, I., Derdak, S., Hutter, B., Eldridge, M.D., Hovig, E., Heisler, L.E., Beck, T.A., Simpson, J.T., Tonon, L. et al. (2015) A comprehensive assessment of somatic mutation detection in cancer using whole-genome sequencing. Nat Commun, 6, 10001.

2. Greaves, M. and Maley, C.C. (2012) Clonal evolution in cancer. Nature, 481, 306–313.

3. Schwarzenbach, H., Hoon, D.S. and Pantel, K. (2011) Cell-free nucleic acids as biomarkers in cancer patients. Nature reviews. Cancer, 11, 426–437.

4. Martincorena, I., Fowler, J.C., Wabik, A., Lawson, A.R.J., Abascal, F., Hall, M.W.J., Cagan, A., Murai, K., Mahbubani, K., Stratton, M.R. et al. (2018) Somatic mutant clones colonize the human esophagus with age. Science (New York, N.Y.), 362, 911–917.

5. Bragg, L.M., Stone, G., Butler, M.K., Hugenholtz, P. and Tyson, G.W. (2013) Shining a light on dark sequencing: characterising errors in Ion Torrent PGM data. PLoS computational biology, 9, e1003031.

6. Pfeiffer, F., Grober, C., Blank, M., Handler, K., Beyer, M., Schultze, J.L. and Mayer, G. (2018) Systematic evaluation of error rates and causes in short samples in next-generation sequencing. Scientific reports, 8, 10950.

7. Fox, E.J., Reid-Bayliss, K.S., Emond, M.J. and Loeb, L.A. (2014) Accuracy of Next Generation Sequencing Platforms. Next generation, sequencing & applications, 1.

8. Xu, C. (2018) A review of somatic single nucleotide variant calling algorithms for next-generation sequencing data. Computational and structural biotechnology journal, 16, 15–24.

9. Gerstung, M., Papaemmanuil, E. and Campbell, P.J. (2014) Subclonal variant calling with multiple samples and prior knowledge. Bioinformatics (Oxford, England), 30, 1198–1204.

10. Martincorena, I., Roshan, A., Gerstung, M., Ellis, P., Van Loo, P., McLaren, S., Wedge, D.C., Fullam, A., Alexandrov, L.B., Tubio, J.M. et al. (2015) Tumor evolution. High burden and pervasive positive selection of somatic mutations in normal human skin. Science (New York, N.Y.), 348, 880–886.

11. Shi, W., Ng, C.K.Y., Lim, R.S., Jiang, T., Kumar, S., Li, X., Wali, V.B., Piscuoglio, S., Gerstein, M.B., Chagpar, A.B. et al. (2018) Reliability of Whole-Exome Sequencing for Assessing Intratumor Genetic Heterogeneity. Cell reports, 25, 1446–1457.

12. Di Tommaso, P., Chatzou, M., Floden, E.W., Barja, P.P., Palumbo, E. and Notredame, C. (2017) Nextflow enables reproducible computational workflows. Nature biotechnology, 35, 316–319.

13. Li, H., Handsaker, B., Wysoker, A., Fennell, T., Ruan, J., Homer, N., Marth, G., Abecasis, G. and Durbin, R. (2009) The Sequence Alignment/Map format and SAMtools. Bioinformatics (Oxford, England), 25, 2078–2079.

14. Aeberhard, W.H., Cantoni, E. and Heritier, S. (2014) Robust inference in the negative binomial regression model with an application to falls data. Biometrics, 70, 920–931.

15. Benjamini, Y. and Hochberg, Y. (1995) Controlling the False Discovery Rate: A Practical and Powerful Approach to Multiple Testing. Journal of the Royal Statistical Society. Series B (Methodological), 57, 289–300.

16. George, J., Lim, J.S., Jang, S.J., Cun, Y., Ozretic, L., Kong, G., Leenders, F., Lu, X., Fernandez-Cuesta, L., Bosco, G. et al. (2015) Comprehensive genomic profiles of small cell lung cancer. Nature, 524, 47–53.

17. Cancer Genome Atlas Research Network. (2012) Comprehensive genomic characterization of squamous cell lung cancers. Nature, 489, 519–525.

18. Ewing, A.D., Houlahan, K.E., Hu, Y., Ellrott, K., Caloian, C., Yamaguchi, T.N., Bare, J.C., P’ng, C., Waggott, D., Sabelnykova, V.Y. et al. (2015) Combining tumor genome simulation with crowdsourcing to benchmark somatic single-nucleotide-variant detection. Nature methods, 12, 623–630.

19. Poplin, R., Ruano-Rubio, V., DePristo, M.A., Fennell, T.J., Carneiro, M.O., Van der Auwera, G.A., Kling, D.E., Gauthier, L.D., Levy-Moonshine, A., Roazen, D. et al. (2018) Scaling accurate genetic variant discovery to tens of thousands of samples. bioRxiv, 201178.

20. Fernandez-Cuesta, L., Perdomo, S., Avogbe, P.H., Leblay, N., Delhomme, T.M., Gaborieau, V., Abedi-Ardekani, B., Chanudet, E., Olivier, M., Zaridze, D. et al. (2016) Identification of Circulating Tumor DNA for the Early Detection of Small-cell Lung Cancer. EBioMedicine, 10, 117–123.

21. Chen, L., Liu, P., Evans, T.C., Jr. and Ettwiller, L.M. (2017) DNA damage is a pervasive cause of sequencing errors, directly confounding variant identification. Science (New York, N.Y.), 355, 752–756.

22. Laehnemann, D., Borkhardt, A. and McHardy, A.C. (2016) Denoising DNA deep sequencing data-high-throughput sequencing errors and their correction. Briefings in bioinformatics, 17, 154–179.

23. Ioannidis, N.M., Rothstein, J.H., Pejaver, V., Middha, S., McDonnell, S.K., Baheti, S., Musolf, A., Li, Q., Holzinger, E., Karyadi, D. et al. (2016) REVEL: An Ensemble Method for Predicting the Pathogenicity of Rare Missense Variants. American journal of human genetics, 99, 877–885.

24. Nong, J., Gong, Y., Guan, Y., Yi, X., Yi, Y., Chang, L., Yang, L., Lv, J., Guo, Z., Jia, H. et al. (2018) Circulating tumor DNA analysis depicts subclonal architecture and genomic evolution of small cell lung cancer. Nat Commun, 9, 3114.

25. LaFramboise, T. (2009) Single nucleotide polymorphism arrays: a decade of biological, computational and technological advances. Nucleic acids research, 37, 4181–4193.

26. Allhoff, M., Schonhuth, A., Martin, M., Costa, I.G., Rahmann, S. and Marschall, T. (2013) Discovering motifs that induce sequencing errors. BMC bioinformatics, 14 Suppl 5, S1.

27. Mose, L.E., Wilkerson, M.D., Hayes, D.N., Perou, C.M. and Parker, J.S. (2014) ABRA: improved coding indel detection via assembly-based realignment. Bioinformatics (Oxford, England), 30, 2813–2815.

28. Kivioja, T., Vaharautio, A., Karlsson, K., Bonke, M., Enge, M., Linnarsson, S. and Taipale, J. (2011) Counting absolute numbers of molecules using unique molecular identifiers. Nature methods, 9, 72–74.

29. Ravasio, V., Ritelli, M., Legati, A. and Giacopuzzi, E. (2018) GARFIELD-NGS: Genomic vARiants FIltering by dEep Learning moDels in NGS. Bioinformatics (Oxford, England), 34, 3038–3040.

30. Boettiger, C. (2015) An introduction to Docker for reproducible research. SIGOPS Oper. Syst. Rev., 49, 71–79.

31. Kurtzer, G.M., Sochat, V. and Bauer, M.W. (2017) Singularity: Scientific containers for mobility of compute. PloS one, 12, e0177459.

## REFERENCES

1. Aeberhard, W.H., Cantoni, E. and Heritier, S. (2014) Robust inference in the negative binomial regression model with an application to falls data. Biometrics, 70, 920–931.

2. Di Tommaso, P., Chatzou, M., Floden, E.W., Barja, P.P., Palumbo, E. and Notredame, C. (2017) Nextflow enables reproducible computational workflows. Nature biotechnology, 35, 316–319.

3. Perkel, J. (2016) Democratic databases: science on GitHub. Nature, 538, 127–128.

4. Merkel, D. (2014) Docker: lightweight Linux containers for consistent development and deployment. Linux J., 2014, 2.

5. Gruning, B., Dale, R., Sjodin, A., Chapman, B.A., Rowe, J., Tomkins-Tinch, C.H., Valieris, R. and Koster, J. (2018) Bioconda: sustainable and comprehensive software distribution for the life sciences. Nature methods, 15, 475–476.

6. Nong, J., Gong, Y., Guan, Y., Yi, X., Yi, Y., Chang, L., Yang, L., Lv, J., Guo, Z., Jia, H. et al. (2018) Circulating tumor DNA analysis depicts subclonal architecture and genomic evolution of small cell lung cancer. Nat Commun, 9, 3114.

7. Martincorena, I., Fowler, J.C., Wabik, A., Lawson, A.R.J., Abascal, F., Hall, M.W.J., Cagan, A., Murai, K., Mahbubani, K., Stratton, M.R. et al. (2018) Somatic mutant clones colonize the human esophagus with age. Science (New York, N.Y.), 362, 911–917.

8. Poplin, R., Ruano-Rubio, V., DePristo, M.A., Fennell, T.J., Carneiro, M.O., Van der Auwera, G.A., Kling, D.E., Gauthier, L.D., Levy-Moonshine, A., Roazen, D. et al. (2018) Scaling accurate genetic variant discovery to tens of thousands of samples. bioRxiv, 201178.

9. Mose, L.E., Wilkerson, M.D., Hayes, D.N., Perou, C.M. and Parker, J.S. (2014) ABRA: improved coding indel detection via assembly-based realignment. Bioinformatics (Oxford, England), 30, 2813–2815.

